# Cognitive resilience despite metabolic dysfunction after adolescent-onset high-fat high-sucrose diet exposure in rats

**DOI:** 10.64898/2026.07.02.736000

**Authors:** Marcia Spoelder, Ilse van Donkelaar, Casper J.H. Wolf, Yami Bright, Sylvia Docq, Anthonieke Middelman, Judith R. Homberg

**Author notes:** Department of Primary and Community Care, Radboud Institute for Medical Innovation, Radboud University Medical Center, Geert Grooteplein Zuid 21, Nijmegen, 6525 GA, The Netherlands.

## Abstract

Adolescence is a sensitive period during which unhealthy diets may shape metabolic health and cognition. Diets high in fat and sugar have been linked to obesity, impaired glucose regulation and hippocampus-dependent deficits, but the exposure duration required to affect cognition remains unclear. This study examined whether adolescent-onset exposure to a high-fat high-sucrose (HFHS) diet induces metabolic dysfunction and impairs object-based memory, spatial working memory and spatial pattern separation in male Long Evans rats.

Rats were assigned to a control or HFHS diet at four weeks of age and remained on this diet into adulthood. Basal blood glucose was assessed monthly and home-cage behaviour using 48-hour LABORAS recordings. Cognitive testing started after 10 weeks of diet exposure, when basal glucose was elevated in HFHS-fed rats. Object displacement and novel object recognition were used in short open-field test settings, whereas touchscreen-based trial-unique nonmatching-to-location testing (TUNL) assessed spatial working memory and pattern separation across repeated operant sessions. Finally, glucose (in)tolerance and tissue weights were measured.

HFHS diet exposure produced a metabolic phenotype, including increased body weight, elevated basal glucose, impaired glucose tolerance and increased liver and gonadal white adipose tissue weights. The diet also altered the general behavioural repertoire, with increased immobility and grooming and reduced rearing. HFHS-fed rats did not differ from controls in object displacement or novel object recognition performance. In the touchscreen task, both groups acquired the task at a comparable rate. Long-delay and spatial separation challenges reduced performance as expected, confirming task sensitivity, but did not reveal diet-related impairments.

These findings show that adolescent-onset HFHS diet exposure induces metabolic dysfunction but does not necessarily produce detectable cognitive impairment when behavioural testing starts after 10 weeks of exposure. Longer exposure or advanced diet-induced inflammatory or neurobiological alterations may be required to reveal cognitive consequences.

## Introduction

The consumption of energy-dense diets rich in saturated fat and sugar contributes to obesity, impaired glucose regulation and other metabolic disturbances. In addition to peripheral metabolic consequences, high-fat and high-sugar diets may affect brain function and cognitive performance, particularly in domains that depend on hippocampal integrity [1–5]. The hippocampus is involved in spatial learning, recognition memory and the discrimination between overlapping spatial representations. These cognitive processes may be vulnerable to diet-induced metabolic, inflammatory and neurobiological changes, although the magnitude and timing of such effects vary across studies [4–6].

Adolescence may represent a particularly sensitive developmental window for diet-induced effects on cognition. During this period, brain regions involved in memory, reward processing and behavioural regulation continue to mature, while dietary habits and metabolic trajectories are also being shaped [6–8]. A systematic review of animal studies concluded that high-fat and high-sugar diet exposure during adolescence often produces stronger or more persistent memory impairments than similar exposure during adulthood [6]. Experimental studies have reported that juvenile or adolescent exposure to high-fat or high-sugar diets can impair hippocampus-dependent memory, alter neurogenesis or synaptic plasticity, and increase inflammatory signalling in the hippocampus [9–13]. More recent work further suggests that obesogenic diet exposure during adolescence can induce circuit-specific alterations in object-based memory through ventral hippocampal projections [14].

However, the relationship between obesogenic diet exposure and cognition is not uniform. Some studies have reported clear impairments in spatial or object-location memory after high-fat or high-sugar diet exposure, whereas others found metabolic, microbiome or neurobiological changes without measurable learning or memory deficits [14–16]. Differences in diet composition, sucrose content, exposure duration, age at exposure, species, sex, strain and behavioural task may all influence whether cognitive effects are detected. Thus, negative or null behavioural findings are informative when the dietary manipulation is effective and when task sensitivity can be demonstrated.

Object displacement and novel object recognition are widely used short behavioural tests to assess object-location and recognition memory in rodents [17,18]. In contrast, touchscreen-based operant tasks allow repeated assessment of cognitive performance across multiple sessions using automated stimulus presentation and response registration. The trial-unique nonmatching-to-location task (TUNL) was developed as a touchscreen test of location memory, spatial working memory and pattern separation [19–21]. In this task, increasing the delay between sample and choice phases increases working memory demand, whereas reducing the spatial separation between response locations increases the demand on spatial discrimination and pattern separation. Recent TUNL and touchscreen pattern separation studies further support the use of these paradigms to study spatial working memory, response strategies, hippocampal-prefrontal mechanisms and dentate gyrus-dependent pattern separation [22–25].

The present study examined whether adolescent-onset exposure to a high-fat high-sucrose (HFHS) diet induces metabolic dysfunction and impairs cognitive performance in male Long Evans rats. We hypothesized that HFHS diet exposure would induce a metabolic phenotype and impair performance in object-based memory, spatial working memory and spatial pattern separation.

## Materials and methods

### Animals, housing and ethical approval

The methods of this study are reported in accordance with the ARRIVE guidelines [26]. The experiment was performed under project licence AVD1030020198164, issued by the Centrale Commissie Dierproeven, The Hague, The Netherlands, for the project ‘*The impact of milk fat globule membrane on cognition in healthy rats and rats exposed to a HFHS diet’*. All procedures were conducted in accordance with Dutch legislation on the use and protection of laboratory animals and were approved locally within Radboud University Medical Center.

A total of 28 male Long Evans rats (*Rattus norvegicus*; RjOrl, Janvier Labs, France) arrived at postnatal day 21, with an approximate body weight of 70 g. Rats were used as the experimental unit. The sample size was based on previous behavioural experiments and the expectation that HFHS diet exposure would induce an approximately 15% disturbance in learning and memory performance. A group size of 13 animals was expected to provide approximately 78% power at an alpha level of 0.05 for a two-group comparison. One additional animal per group was included to account for potential failure to acquire the touchscreen task.

Upon arrival, rats were housed in the animal facility of Radboud University Medical Center under a reversed light–dark cycle to facilitate behavioural testing during the active phase. Rats were socially housed in pairs in Eurostandard type IV cages for most of the experiment. Individual housing in type III cages was limited to predefined periods required for individual food and water intake measurements and for LABORAS recordings. During LABORAS measurements, animals were individually housed for 48 h. Water was available ad libitum throughout the experiment. Food was available ad libitum except during cognitive testing periods, when food was removed 3–4 h before touchscreen sessions to increase task engagement. Immediately after testing, rats again received ad libitum access to their assigned diet.

All rats were individually identified by tail colour marking. Body weight and general health were monitored throughout the experiment and measured at least twice weekly.

### Study design and dietary intervention

Baseline blood glucose was measured for the first time after either two- or four-days following arrival. After the blood glucose measurement, rats were allocated to one of two dietary groups: a control diet group (n = 14) and a high-fat high-sucrose diet group (HFHS; n = 14). The HFHS group received the purified HFHS diet D12451, containing high (45%) levels of saturated fat (with lard and soybean oil) and sucrose, containing 4.615 kcal/kg, 10mm pellets, red color added, gamma-irradiated 25 kGy (Sniff diets, Bio-Services B.V., Schaijk, the Netherlands). The control group received the matched purified control diet D12450K, containing 3.630 kcal/kg, 10mm pellets, gamma-irradiated 25 kGy. Both diets were provided in pelleted form in the food hopper of the home cage.

The general experimental timeline consisted of baseline metabolic assessment after habituation, followed by dietary induction during adolescence and cognitive testing during early and general adulthood while animals remained on their assigned diets. Body weight was measured repeatedly throughout the study. Blood glucose was assessed once per month, and 48h LABORAS recordings were performed every month as well. Object displacement and novel object recognition testing were performed first after 10 weeks on diet exposure and a second time after about 24 weeks on diet exposure. TUNL training and testing were performed in the 14 weeks between the two NORT sessions.

### Randomisation and blinding

Animals were assigned to the control or HFHS diet groups, and cage positions within the housing room and the LABORAS room were alternated between dietary groups to avoid systematic environmental bias due to rack position or location in the room. The order in which animals were exposed to blood draw, or the timing of behavioral testing, was randomized.

Complete blinding during all procedures was not feasible because the HFHS diet was visibly more fatty and differed in color (red color) from the control diet. In addition, the experimenters monitored food intake and handled animals from cages in which the assigned diet was visible. Behavioural testing itself was therefore not fully blinded, but automated data acquisition in the touchscreen chambers reduced observer-dependent outcome scoring for the TUNL task. Where feasible, blinding was applied during sample collection; for example, during blood collection one person removed the animal from the home cage while the person collecting the blood sample remained unaware of group allocation.

### Inclusion and exclusion criteria

All animals that entered the experiment were included in longitudinal metabolic and general behavioural analyses unless data were missing because of technical failure or health-related exclusion. For the TUNL task, animals were required to acquire the task before progressing to the subsequent test stages. Animals that did not reach this criterion within the available training period were excluded from subsequent TUNL test-stage analyses but remained part of the study population for endpoints for which valid data were available.

### General behavioural assessment using LABORAS

General home-cage behaviour was assessed repeatedly using the Laboratory Animal Behavior Observation Registration and Analysis System (LABORAS; Metris B.V., Hoofddorp, The Netherlands). LABORAS is an automated system that detects vibrations generated by the animal’s movements and classifies these into behavioural parameters such as locomotion, resting, grooming, rearing, eating and drinking [27]. During these sessions, animals had access to food and water in the cage rack.

### Blood glucose, glucose tolerance testing and liver and adipose tissue weights

Blood glucose was measured repeatedly as an indicator of metabolic homeostasis. Blood samples were collected once per month. Before blood sampling, food was removed for 5 h. To facilitate blood collection, animals were briefly placed under a heat lamp to dilate the tail veins. Blood was collected from the tail vein using a refined approach with a needle and capillary tube where possible, thereby avoiding tail incision unless necessary.

The oral glucose tolerance test was performed during the final phase of the experiment, after completion of the main cognitive TUNL test period. After 12 h of food restriction, baseline blood glucose was measured (t = 0), after which rats received an oral load of 50% D-glucose at a dose of 2 g/kg body weight. Blood glucose was subsequently measured at 15, 30, 60, 90 and 120 min after glucose administration.

At the end of the experiment, rats were euthanized by exposure to CO₂. Immediately after euthanasia, the liver, gonadal white adipose tissue and mesenteric white adipose tissue were dissected and weighed.

### Object displacement (OD) and novel object recognition task (NORT)

Spatial and recognition memory were assessed using the object displacement task and the novel object recognition task, respectively. Both tasks exploit the innate tendency of rats to preferentially explore a displaced or novel object over a familiar object [17,18,28]. The general set-up was comparable across both tasks and consisted of habituation, training and testing phases.

During the first habituation session, rats were placed in the empty arena for 30 min without objects or wall cues. During subsequent habituation sessions, rats were allowed to explore the arena for 10 min. On the training day, two identical objects were placed in the arena and rats were allowed to explore for 10 min. Visual cues were placed on the walls of the arena, and one three-dimensional cue was used to facilitate spatial orientation. On the test day of the object displacement task, one of the familiar objects was moved to a new location. On the test day of the novel object recognition task, one familiar object was replaced by a novel object. Different objects were used across tasks and across repeated test occasions.

The object displacement and novel object recognition tasks were performed twice: once during early adulthood (10 weeks old) and once at the end of the experiment in general adulthood (24 weeks old). For the second test occasion, new objects and cues were used. The second NORT procedure included one habituation day with two sessions: first, rats were placed in the arena without cues or objects for 5 min with their cage mate; second, rats were individually allowed to explore two objects placed in the center of the arena for 5 min. The primary outcomes were the relative exploration of the displaced object compared with the non-displaced object in the object displacement task, and the relative exploration of the novel object compared with the familiar object in the novel object recognition task. Total exploration time was assessed to evaluate potential group differences in general exploratory behaviour.

### Trial-unique nonmatching-to-location task

Spatial working memory and pattern separation were assessed using the trial-unique nonmatching-to-location task (TUNL), performed in sound-attenuated Bussey–Saksida touchscreen operant chambers for rats (Campden Instruments, Loughborough, UK) [19–21,29,30]. Each chamber contained a touchscreen on one side and a food pellet dispenser with a magazine on the opposite side. A polycarbonate mask, dividing the touchscreen in 15 equal squares, was placed in front of the touch screen from the ‘Must touch’-phase onwards. Stimulus presentation, reward delivery and response registration were controlled using Whisker and ABET II software for Operant Control [31]. The food reinforcement pellets used were 45 mg grain-based precision pellets purchased from TestDiet (Purina Mills LLC, Richmond, IN, USA; Catalog No. 5TUM).

Rats were trained according to the learning schedule described in Table 1. Rats had to reach the prespecified criterion before advancing to the next stage. All training sessions before the TUNL sessions lasted 30 minutes. In the TUNL task, each trial consisted of a sample phase and a choice phase. During the sample phase, one location on the touchscreen was illuminated, which the rat needed to press to get a food reward in 33% of trials. After a head entry into the food magazine and subsequently one second delay, two locations were illuminated during the choice phase: the previously presented sample location and a new location. The rat had to select the new, nonmatching location to receive a food reward.

**Table 1.** Different stages of cognitive testing and training.

| **TUNL training** |  |  |
| --- | --- | --- |
| **Learning stage** | **Criteria** | **Description** |
| **Habituation** | Complete one session | The rat was provided with 25 pellets at variable time intervals. |
| **Autoshaping** | All 25 pellets eaten | To learn the association between the stimulus and the reward, the rat received a reward regardless of whether it touched the active stimulus and would receive the reward |
|  |  | immediately when it touched the active stimulus. |
| **Must touch** | Two sessions 25 correct responses and all 25 pellets eaten | The rat had to touch the active stimulus to receive a reward, otherwise no reward was given. Across three must-touch stages, the active response area was progressively reduced until only one square remained. |
| **Punish incorrect** | At least 70% correct responses for two sessions | When the active stimulus was touched, the rat received a food reward. When an inactive stimulus was touched, the rat received negative feedback (house light on) and no food reward was provided. |
| **TUNL** | At least 15 sessions and at least 70% correct responses on average for three subsequent sessions and less than 15% variability in correct responding within these three sessions | The rat had to touch a single active stimulus. After a delay of one second, two squares are active, and a response in the new lit location is required to receive a reward. |
| **TUNL long delay** | 2 sessions | Increasing the delay between sample and choice phase from 1 s to 6 s |
| **TUNL** | 2 sessions |  |
| **TUNL long delay** | 2 sessions | Increasing the delay between sample and choice phase from 1 s to 6 s |
| **TUNL intermediate separation** | 3 sessions | Varying the spatial separation between the sample and choice locations per trial with large and intermediate separation |
| **TUNL small separation** | 3 sessions | Varying the spatial separation between the sample and choice locations with large, intermediate and small separation |

Incorrect responses and omissions were followed by negative feedback comparable to the punish-incorrect stage. TUNL sessions lasted 45 min each. Rats were required to complete at least 15 TUNL A sessions and to achieve at least 70% correct responses on average for three subsequent sessions and less than 15% variability in correct responding within these three sessions, to meet the TUNL acquisition criterion.

During the initial TUNL A acquisition stage, the delay between sample and choice phase was 1 s. After acquisition, task difficulty was manipulated by increasing the delay between sample and choice phase from 1 s to 6 s and by varying the spatial separation between the sample and choice locations. Large-separation trials were considered easier, medium-separation trials intermediate, and small-separation trials most difficult because the two active response locations were adjacent or near-adjacent. The primary TUNL outcome was percentage correct responses, calculated as the number of correct choice responses divided by the total number of completed choice trials.

### Statistical analysis

The statistical analysis was designed to compare the control and HFHS groups across metabolic, general behavioural and cognitive outcomes, using the individual rat as the experimental unit. Analyses were performed using IBM SPSS Statistics, version 31.

Descriptive statistics in the result text is presented as mean ± SD. Graphical representations were generated using GraphPad Prism version 11 (GraphPad Software, Boston, Massachusetts, USA). Data are visualized as mean ± SEM, with or without individual data points.

Repeatedly measured outcomes, including body weight, glucose levels and LABORAS-derived behavioural parameters, were analyzed using general linear models for repeated measures. Within the HFHS-fed group, one rat died after the second blood draw procedure, and another rat was sacrificed due to a broken foot after 5 months in the experiment. One control rat died, without a defined reason after 4 months. Hence, these animals were not included within the statistical analyses. Diet group was included as between-subject factor and time or session as within-subject factor. Homogeneity of variance was assessed before parametric group comparisons. When repeated-measures assumptions were not met, Greenhouse–Geisser correction will be applied. Normality was assessed by visual inspection and, where appropriate, applied with Shapiro–Wilk tests. Group-by-time interactions were used to determine whether developmental or diet-induced trajectories differ between groups. Post hoc pairwise comparisons were only performed following significant main effects or interactions. Pairwise comparisons with a Bonferroni correction were used for follow-up comparisons at individual time points. For the TUNL task, percentage correct responses were analyzed across acquisition and test stages. Training performance was analyzed by comparing the number of sessions required to reach criterion and percentage correct responses across sessions with independent samples two-sided t-tests. Test-stage performance (challenges) was analyzed using repeated-measures models with group as between-subject factor and session, delay and/or separation as within-subject factors where applicable.

For the object displacement task and the novel object recognition task, the discrimination index (DI) was used to assess memory performance. This was calculated as the difference in time exploring the novel object location or new object and stable location divided or same object by the total exploration time. This results in a score ranging from −1 (preference for the stable location/same object) to +1 (preference for the moving object location/new object). A score of 0 indicates no preference for either object location. Total exploration time was also analyzed to distinguish memory performance from differences in general exploration. The DI and the total exploration time was analyzed between groups with independent samples T-tests.

The area under the curve for the glucose tolerance test, the tissue weights were tested with independent samples one-sided t-tests.

## Results

### Body weight and blood glucose development

Body weight did not differ between groups at baseline (week 0), before the start of the dietary intervention (HFHS: 73.3 ± 7.8 g; control: 70.5 ± 8.5 g; p = 0.367). During the dietary intervention, HFHS-fed rats showed a progressively higher body weight compared with control-fed rats (Fgroup*week_(27, 621)_ = 6.36, p < 0.001; Fgroup _(1, 23)_ = 8.30, p = 0.008). This difference was already apparent after 1 week of diet exposure (HFHS: 111.3 ± 10.0 g; control: 99.7 ± 13.7 g; p = 0.019) and persisted throughout the experiment. At week 27, HFHS-fed rats weighed 11% more; 572 ± 48 g compared with 517 ± 25 g in the control group (p = 0.001) (Fig 1A).

**Figure 1.**
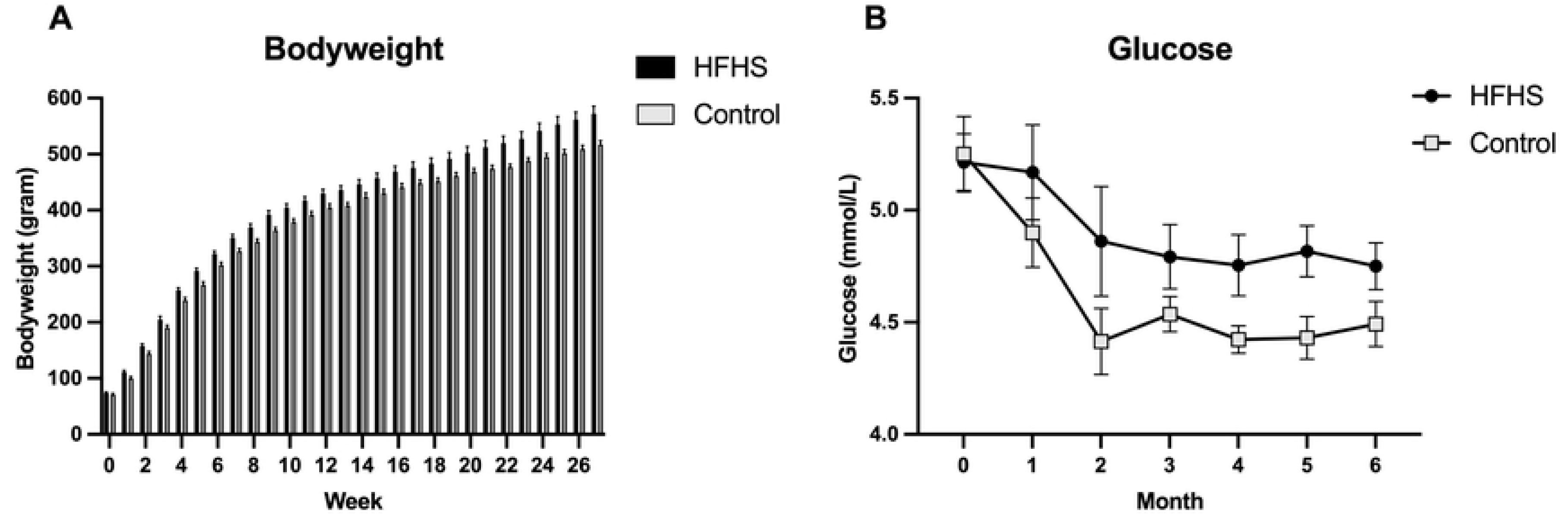
Body weight and basal blood glucose levels in control and HFHS-fed rats. (A) Body weight development during the 27 weeks of diet exposure. (B) Basal blood glucose levels measured monthly during the experiment.

Baseline blood glucose levels did not differ between groups before the start of the dietary intervention (week 0). During the experiment, HFHS-fed rats developed higher basal glucose levels than control-fed rats across the repeated monthly measurements (Fgroup _(1, 23)_ = 7.83, p = 0.010), but there was no significant group × time interaction (Fgroup*month _(6, 138)_ = 0.30, p = 0.834) (Fig 1B).

### General behavior over 48 sessions in LABORAS

The 48h sessions of measuring the general behavioral repertoire was monthly repeated and consisted of seven subsequent sessions. The duration spent on locomotor activity decreased once the rats aged and was indifferent between groups (Fmonth _(4, 99)_ = 15.24, p < 0.001; Fgroup*month _(4, 99)_ = 1.17, p = 0.330; Fgroup _(1, 23)_ = 0.12, p = 0.735) (Fig 2A). Instead, immobility increased over time and was higher for HFHS-fed rats (Fmonth _(4, 94)_ = 208.08, p < 0.001; Fgroup*month _(4, 94)_ = 1.22, p = 0.308; Fgroup _(1, 23)_ = 4.88, p = 0.037) (Fig 2B). Rearing and grooming behaviours decreased over time, and HFHS-fed rats spent less time on rearing (Fig 2C), but more time on grooming (Fig 2D) (rearing: Fmonth _(3, 77)_ = 24.29, p < 0.001; Fgroup*month _(3, 77)_ = 0.97, p = 0.419; Fgroup _(1, 23)_ = 9.78, p = 0.005; grooming (Fmonth _(2, 41)_ = 67.64, p < 0.001; Fgroup*month _(4, 41)_ = 1.26, p = 0.219; Fgroup _(1, 23)_ = 8.36, p = 0.008).

**Figure 2.**
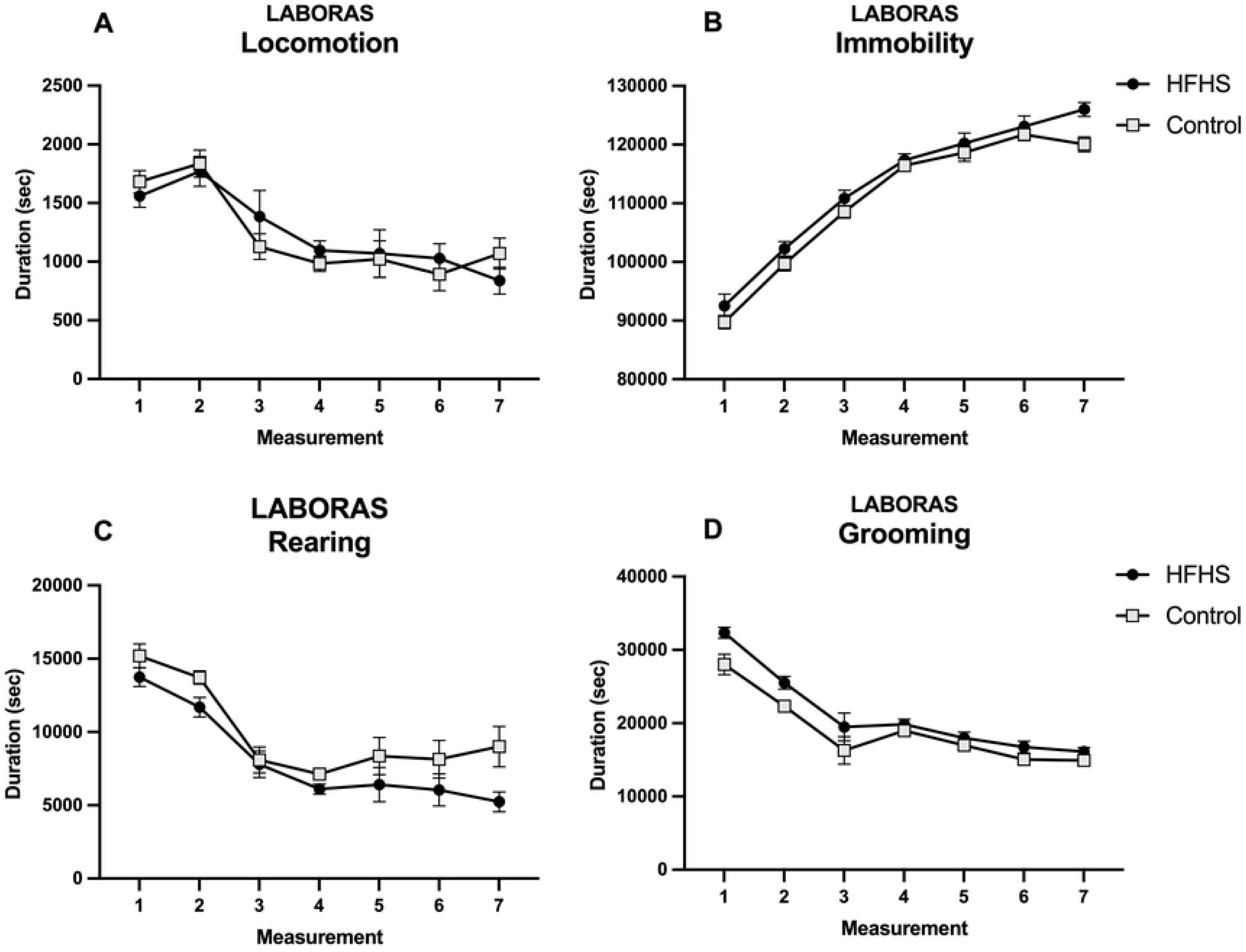
General behavioural repertoire in control and HFHS-fed rats. General behaviour was assessed during repeated 48h LABORAS sessions. (A) Locomotor activity. (B) Immobility. (C) Rearing. (D) Grooming.

The duration spent on drinking water was higher for HFHS-fed rats (Fig 3A), whereas the actual amount of consumed water (corrected for bodyweight) was lower for HFHS-fed rats (Fig 3C). (duration: Fmonth _(2, 55)_ = 14.40, p < 0.001; Fgroup*month _(2, 55)_ = 1.54, p = 0.220; Fgroup _(1, 23)_ = 6.31, p = 0.019; consumed ml: Fmonth _(2, 45)_ = 169.04, p < 0.001; Fgroup*month _(3, 70)_ = 0.45, p = 0.638; Fgroup _(1, 23)_ = 5.91, p = 0.023). The duration spent on eating (Fig 3B) and the actual amount of consumed food was lower for HFHS-fed rats (Fig 3D) (duration: Fmonth _(2, 39)_ = 7.72, p = 0.002; Fgroup*month _(2, 55)_ = 4.87, p = 0.017; Fgroup _(1, 23)_ = 57.28, p < 0.001; consumed food: Fmonth _(2, 36)_ = 145.74, p < 0.001; Fgroup*month _(2, 36)_ = 2.07, p = 0.150; Fgroup _(1, 23)_ = 39.73, p < 0.001). Actual consumed kcal food corrected for BW decreased over time and was not different between groups (Fmonth _(2, 38)_ = 155.51, p < 0.001; Fgroup*month _(2, 38)_ = 0.83, p = 0.423; Fgroup _(1, 23)_ = 3.59, p = 0.071).

**Figure 3.**
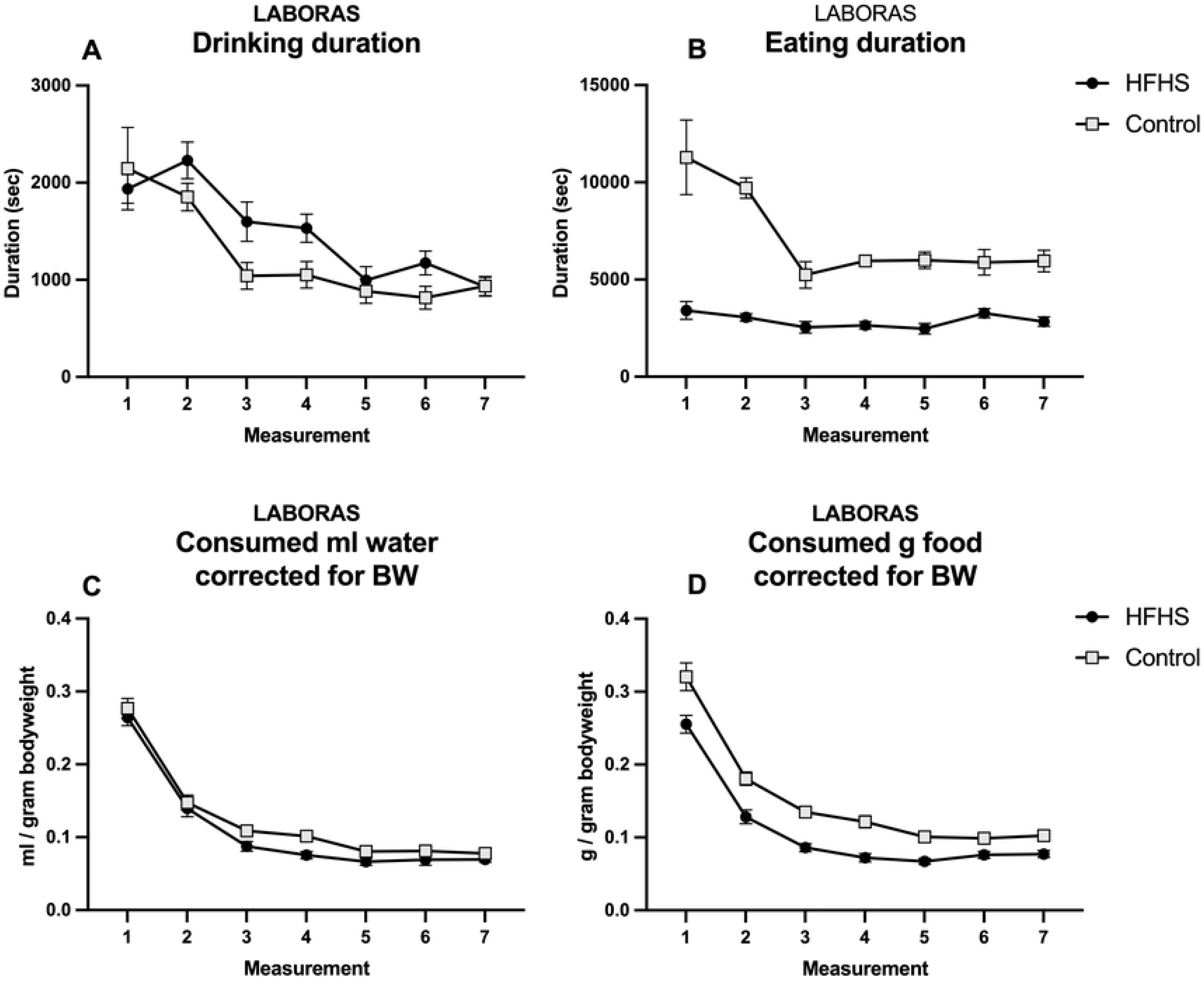
Drinking and eating behaviour in control and HFHS-fed rats. Drinking and eating behaviour were assessed during repeated 48h LABORAS sessions. (A) Duration spent drinking. (B) Duration spent eating. (C) Body weight-corrected water intake. (D) Body weight-corrected food intake.

### Glucose tolerance test

Upon 26 weeks of exposure to the diets, both groups were challenged with the glucose tolerance test. The glucose levels were highest directly after 15 minutes and sustained high in the HFHS-fed rats till 90 minutes after oral gavage, whereas control-fed rats directly reduced their glucose levels (Fgroup*time _(5, 78)_ = 3.75, p = 0.011; Fgroup _(1, 23)_ = 39.57, p < 0.001) (Fig 4A). The area under the curve (AUC) was higher in the HFHS-fed rats (Tgroup _(23)_ = 6.67, p < 0.001) (Fig 4B).

**Figure 4.**
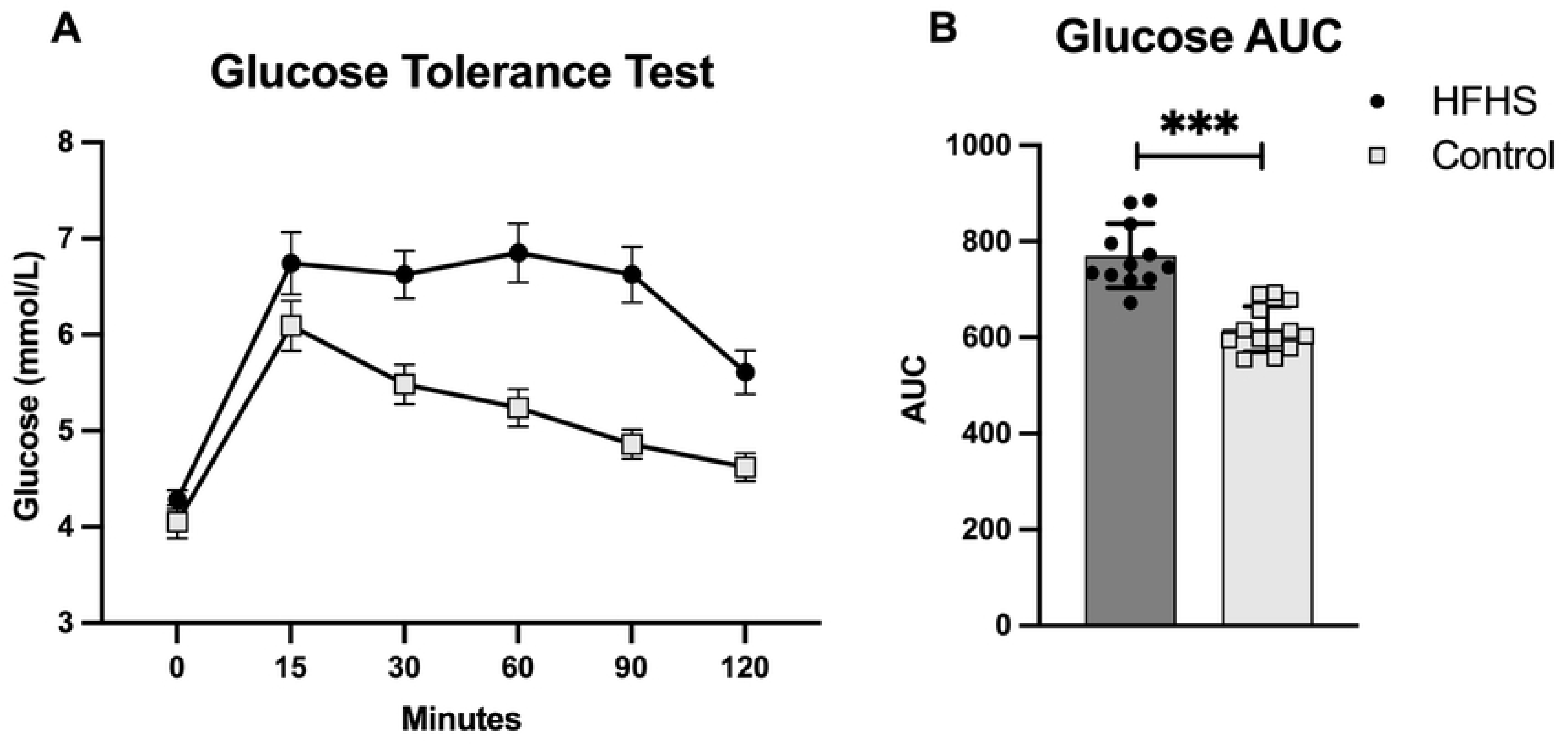
Glucose tolerance in control and HFHS-fed rats. (A) Blood glucose levels during the oral glucose tolerance test after 26 weeks of diet exposure. (B) Area under the curve of the oral glucose tolerance test.

### Liver and white adipose tissue weights

The measured liver weight (Fig 5A) and gonadal white adipose tissue weight (Fig 5B) was higher in the HFHS-fed rats (Tgroup _(13)_ = 2.59, p = 0.011; Tgroup _(22)_ = 5.55, p < 0.001, respectively), but did not reached significance for the mesenteric white adipose tissue (Tgroup _(23)_ = 1.52, p = 0.072) (Fig 5C).

**Figure 5.**
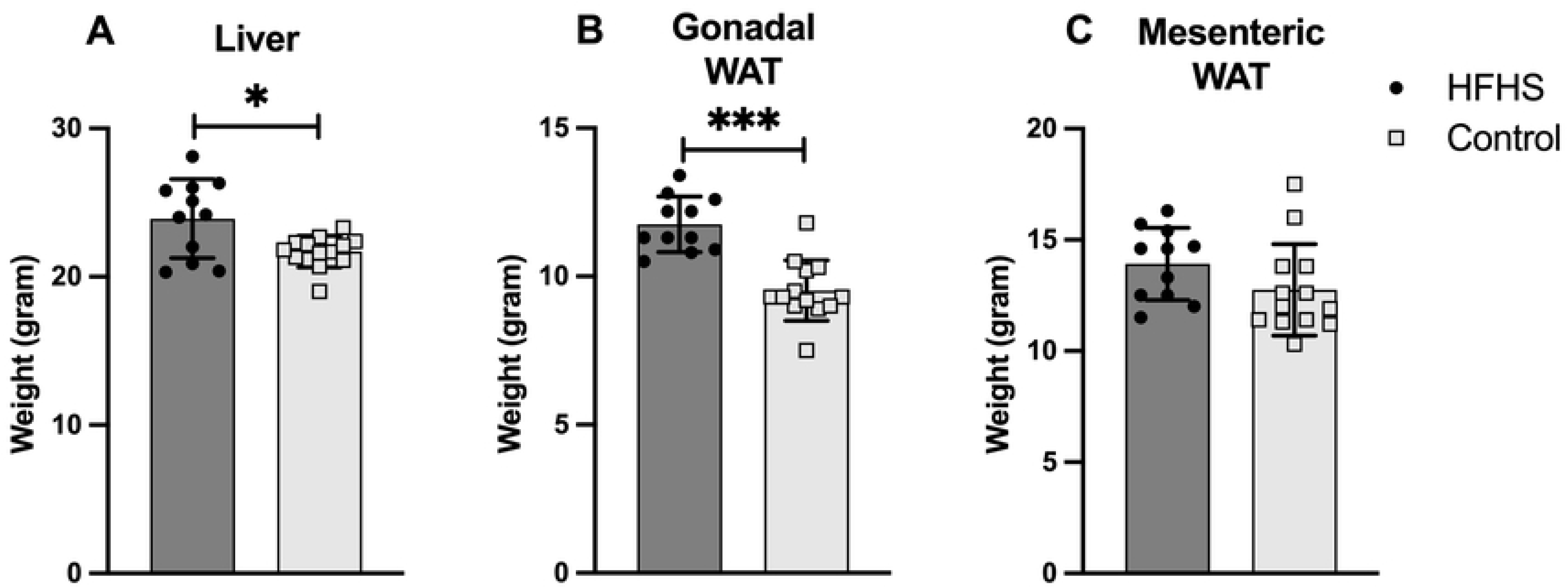
Liver and white adipose tissue weights in control and HFHS-fed rats. Tissue weights were measured at the end of the experiment. (A) Liver weight. (B) Gonadal white adipose tissue weight. (C) Mesenteric white adipose tissue weight.

### Object displacement (OD) and novel object recognition task (NORT)

After 10 weeks of exposure to the diets, when rats were 14 weeks old, the discrimination index (DI) for the new location (OD) or the novel object (NORT) was not different between HFHS-fed rats and control rats (Tgroup _(25)_ = -0.71, p = 0.487; Tgroup _(25)_ = 0.91, p = 0.371, respectively). After 24 weeks of exposure to the diets, the NORT was repeated but no group difference in DI was observed (Tgroup _(23)_ = -0.37, p = 0.716) (Fig 6A). The total exploration time during the OD, and NORT after 10 or 24 weeks on diet exposure was comparable in the two groups (Tgroup _(25)_ = -0.48, p = 0.633; Tgroup _(25)_ = -0.99, p = 0.332; Tgroup _(23)_ = 0.78, p = 0.445, respectively) (Fig 6A).

**Figure 6.**
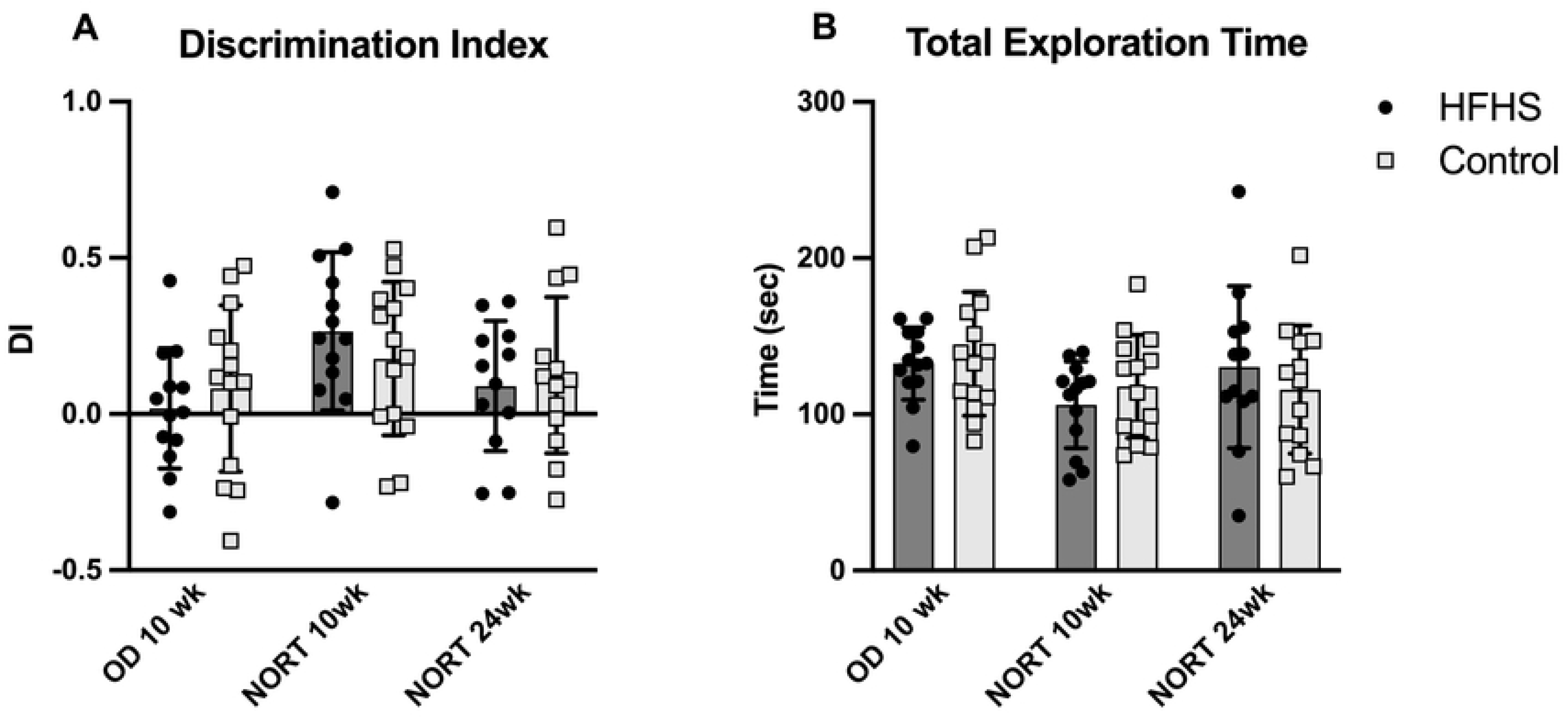
Object displacement and novel object recognition performance in control and HFHS-fed rats. (A) Discrimination index (DI) in the object displacement (OD) task after 10 weeks of diet exposure, the novel object recognition task (NORT) after 10 weeks of diet exposure, and the repeated NORT after approximately 24 weeks of diet exposure. (B) Total exploration time during the corresponding test sessions.

### TUNL Training and test

The number of training sessions during autoshaping (2 ± 0.4 vs 2 ± 0.8), must-touch (7 ± 1.2 vs 8 ± 1.9), and punish incorrect (5 ± 1.5 vs 6 ± 2.2), needed to reach criterion were comparable between HFHS-fed and control-fed rats (Tgroup _(17)_ = -1.29, p = 0.215; Tgroup _(24)_ = -0.86, p = 0.396; Tgroup _(23)_ = -0.91, p = 0.374, respectively).

The groups showed a similar increase in performance during the initial 15 training sessions of the TUNL (Fsession _(14, 322)_ = 4.48, p < 0.001; Fgroup*session _(14, 332)_ = 0.76, p = 0.711; Fgroup _(1, 23)_ = 0.01, p = 0.921) (Fig 7A). Groups did not differ in their average performance during the last three training sessions before the rat progressed to the task difficulty challenges (Tgroup _(16)_ = -0.34, p = 0.740) (Fig 7B) and there was no difference between groups in the number of sessions required to reach the TUNLA criterion (Tgroup _(16)_ = 0.65, p = 0.527) (Fig 7C). Four rats in the control-fed group and three rats in the HFHS-fed group did not reach the TUNLA training criterion and were therefore not progressed into the subsequent challenge sessions. This resulted in 9 vs 9 rats/group to continue in the subsequent challenge sessions.

**Figure 7.**
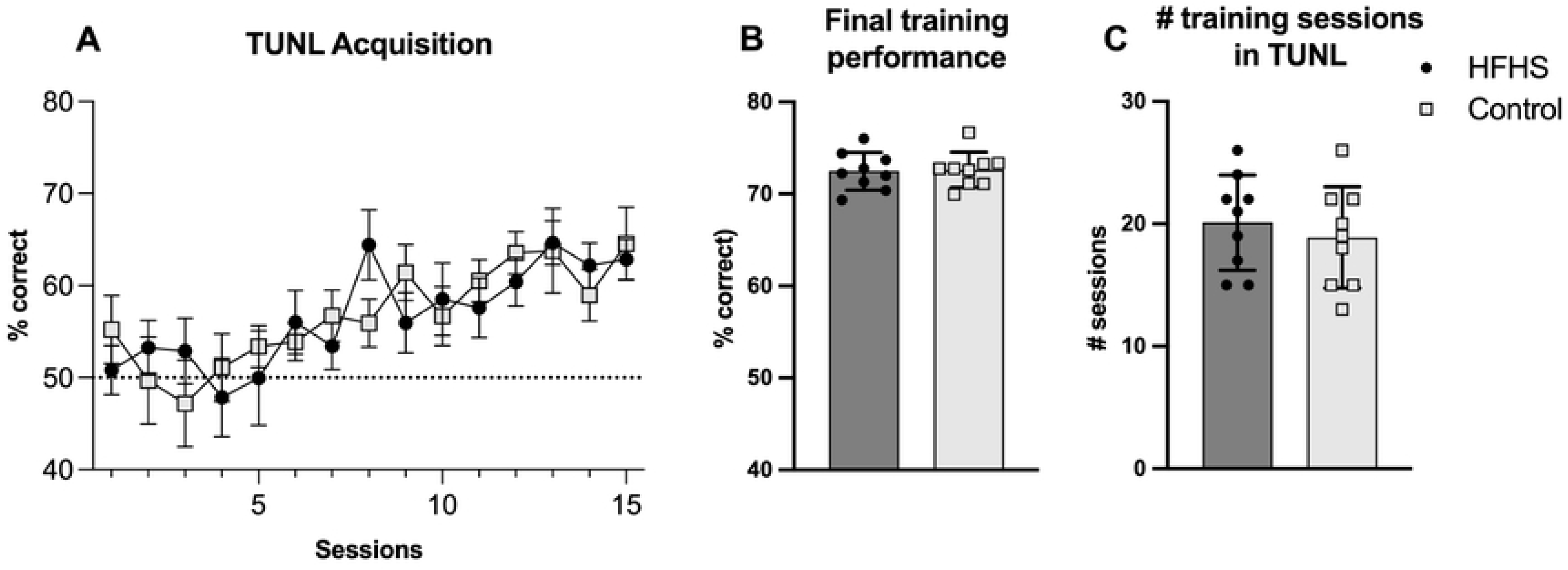
Acquisition of the TUNL task in control and HFHS-fed rats. (A) Percentage of correct responses during the first 15 TUNL acquisition sessions. (B) Average percentage of correct responses during the final three acquisition sessions before progression to the challenge sessions. (C) Number of sessions required to reach the TUNL acquisition criterion.

### TUNL challenge sessions

#### Long delay

The long delay (6 sec) challenge significantly reduced performance compared to the standard short delay (1 sec) between the sample and choice trial within the TUNL task (Fchallenge_(1, 16)_ = 260.61, p < 0.001). The challenge negatively influenced the performance of both groups in a similar manner and did not result in a main group difference (Fgroup*challenge _(1, 16)_ = 0.65, p = 0.433; Fgroup _(1, 16)_ = 0.50, p = 0.491) (Fig 8A). During the long delay challenge sessions, both groups gained less rewards (Fchallenge_(1, 16)_ = 75.12, p < 0.001; Fgroup*challenge _(1, 16)_ = 0.81, p = 0.382; Fgroup _(1, 16)_ = 4.21, p = 0.057), whereas the number of completed trials remained comparable (Fchallenge_(1, 16)_ = 3.08, p = 0.098; Fgroup*challenge _(1, 16)_ = 0.74, p = 0.403; Fgroup _(1, 16)_ = 3.09, p = 0.098).

**Figure 8.**
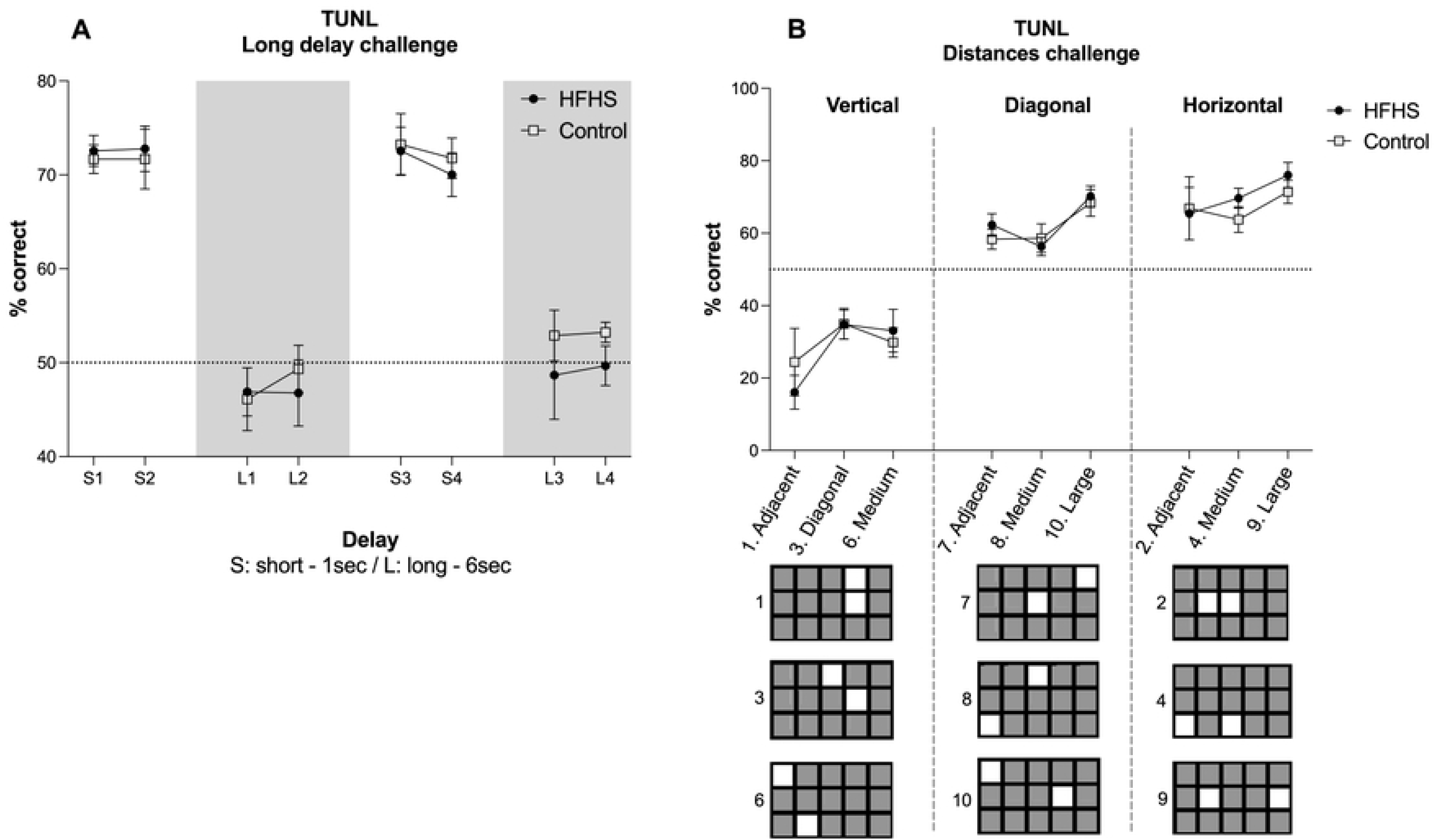
TUNL long-delay and spatial separation challenges in control and HFHS-fed rats. (A) Percentage of correct responses during the standard short-delay and long-delay challenge sessions. (B) Percentage of correct responses across the nine spatial configurations of the separation challenge.

#### Distances – pattern separation

The nine different pattern groups resulted in a significant difference in performance, depending on the pattern group (Fdistances_(3, 49)_ = 30.33, p < 0.001), but independent of the HFHS or the control diet (Fgroup*session _(3, 49)_ = 0.45, p = 0.733; Fgroup _(1, 15)_ = 0.13, p = 0.722) (Fig 8B). Subsequent pairwise comparisons showed that the performance in the three vertical patterns was significantly lower compared to all other six diagonal or horizontal patterns (p<0.001 to p=0.016). The performance on the horizontal sessions with a large separation was significantly higher compared to both the adjacent (p<0.001) and the medium (p<0.019) diagonal separation. Within the diagonal patterns, the medium separation differed also significantly from the large separation (p=0.018).

After grouping the three types of distances into vertical, diagonal and horizontal, the performance within the three pattern types were significantly different (Fdistances_(2, 30)_ = 77.27, p < 0.001), without a significant effect of group (Fgroup*session _(2, 30)_ = 0.26, p = 0.777; Fgroup _(1, 15)_ = 0.45, p = 0.515). Subsequent pairwise comparisons showed that the performance in the vertical patterns was significantly lower compared to the diagonal (p<0.001) and horizontal patterns (p<0.001), and also the diagonal and horizontal patterns differed significantly from each other (p<0.044) (Fig 9A-B).

**Figure 9.**
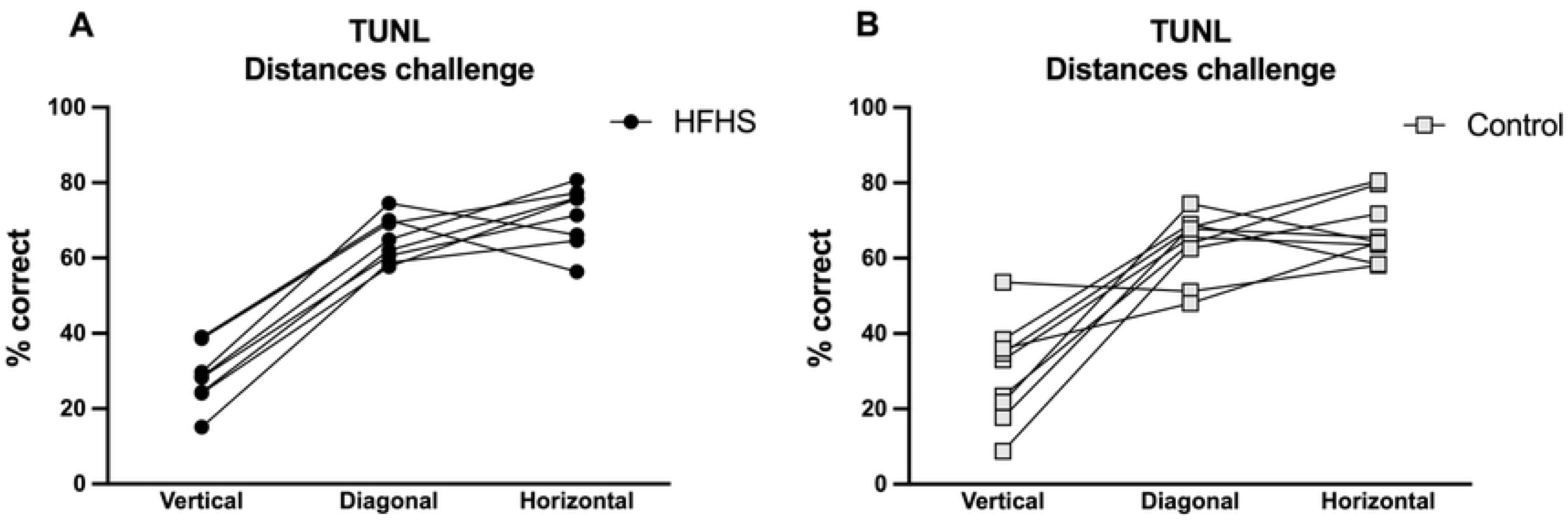
TUNL performance grouped by spatial configuration type in control and HFHS-fed rats. Percentage of correct responses grouped by vertical, diagonal and horizontal spatial configurations for HFHS-fed rats (A) and control-fed rats (B).

## Discussion

In this study, adolescent-onset exposure to a HFHS diet induced a clear metabolic phenotype in male Long Evans rats, but did not lead to detectable impairments in the cognitive tasks used here. Compared with control-fed rats, HFHS-fed rats gained more body weight, developed higher basal glucose levels, showed impaired glucose tolerance and had increased liver and gonadal white adipose tissue weights. The diet also altered aspects of the general behavioural repertoire, with increased immobility and grooming and reduced rearing. However, these metabolic and behavioural changes were not accompanied by impairments in object displacement, novel object recognition, TUNL acquisition, delayed spatial working memory or spatial pattern separation. Importantly, the TUNL challenge sessions reduced performance as expected, indicating that the task manipulations increased cognitive demand.

### Absence of detectable cognitive impairment despite metabolic dysfunction

The central finding of this study is that metabolic dysfunction did not translate into detectable cognitive impairment under the present testing conditions. This contrasts with several studies showing that high-fat or high-sugar diet exposure can impair hippocampus-dependent memory, particularly when exposure occurs during adolescence [6,9–14]. In particular, juvenile or adolescent high-fat diet exposure has been associated with impaired relational memory, object-location memory, reduced hippocampal neurogenesis, altered synaptic plasticity and hippocampal inflammation [9–12]. More recent work has shown that adolescent obesogenic diet exposure can induce circuit-specific object-based memory deficits through ventral hippocampal pathways [14].

At the same time, the present results are consistent with studies showing that diet-induced metabolic or biological changes do not always coincide with measurable impairments in learning or memory. Deshpande et al. reported that high-fat diet exposure altered gut microbiota, but did not affect spatial working memory or microgliosis in early middle-aged Sprague Dawley rats [15]. Similarly, Leyh et al. found that long-term diet-induced obesity in mice did not necessarily lead to learning or spatial memory deficits, despite general health effects and hypothalamic microglial changes [16]. In our previous work in Zucker Diabetic Fatty rats, early diabetic and inflammatory features were also not accompanied by cognitive decline during adolescence [32]. Together, these findings support the view that the relationship between obesogenic diet exposure and cognition depends on diet composition, exposure duration, age at exposure, species or strain, sex, task demands and the neurobiological state at the time of testing.

### Timing of behavioural testing and possible delayed neurobiological consequences

One possible explanation for the absence of detectable cognitive impairment is the timing of behavioural testing relative to the development of diet-induced neurobiological alterations. In the present study, cognitive testing started after 10 weeks of diet exposure, when basal glucose levels were already elevated in HFHS-fed rats. However, elevated glucose and increased body weight may precede the inflammatory, synaptic or hippocampal alterations required to produce robust behavioural deficits. A time-response study in a fructose-induced insulin resistance model showed that insulin resistance appeared from the seventh week of exposure, whereas cognitive dysfunction emerged only after 20 weeks [33]. A recent study on the chronological dynamics of high-fat diet-induced inflammation also suggests that inflammatory responses may progress across peripheral tissues before later involving brain tissue [34].

The timing of individual tasks may also be relevant. Object displacement and the first novel object recognition task were performed relatively early, after approximately 10 weeks of diet exposure. The repeated novel object recognition task and the later phases of TUNL testing occurred after longer exposure, but by that time the animals had already undergone extended training and repeated cognitive testing. It remains possible that stronger cognitive effects would have emerged if all cognitive testing had started only after more prolonged diet exposure, or if behavioural testing had been combined with direct measures of hippocampal inflammation, synaptic plasticity, insulin signalling or neurogenesis. Studies reporting diet-induced cognitive impairments often implicate hippocampal plasticity, glucocorticoid signalling, neuroinflammation or blood-brain barrier changes as potential mechanisms [10,12,35].

### Interpretation of the TUNL findings

The TUNL task was included to assess spatial working memory and pattern separation across repeated operant sessions. This task requires animals to select a novel location after a delay and allows task difficulty to be manipulated by increasing the delay or reducing the spatial separation between locations [19–21]. In the present study, both groups acquired the task at a comparable rate, and their performance during the final acquisition sessions did not differ. The long-delay challenge reduced performance in both groups, consistent with increased working memory demand. Spatial separation challenges also produced clear distance-dependent changes in accuracy, consistent with increased demands on spatial discrimination and pattern separation. However, neither manipulation revealed a diet-related deficit.

Recent TUNL studies further support both the sensitivity and the complexity of this task. Pharmacological and electrophysiological studies show that TUNL performance can be altered by N-methyl-D-aspartate receptor and dopaminergic manipulations and is linked to prefrontal-hippocampal activity and task engagement [36,37]. Computational and intervention studies further indicate that TUNL performance reflects an integrated behavioural outcome, including working memory components, response biases and condition-specific sensitivity [37,38]. In addition, touchscreen-based spatial separation paradigms recruit dentate gyrus activity under high cognitive demand [25]. Thus, the present challenge effects support increased cognitive demand, while the absence of diet effects should be interpreted within the multidimensional nature of TUNL performance.

### Strengths and limitations

A strength of this study is the longitudinal design, in which diet exposure started during adolescence and continued into adulthood. The study combined repeated metabolic assessments, automated home-cage behavioural monitoring and multiple cognitive tasks. The inclusion of both short open-field memory tests and repeated touchscreen-operant testing allowed assessment of object-location memory, recognition memory, spatial working memory and spatial pattern separation. In addition, the TUNL task included performance challenges that confirmed sensitivity to delay and spatial configuration.

Several limitations should also be considered. First, only male rats were included, so the findings cannot be generalized to females. Second, cognitive testing started after 10 weeks of diet exposure. Although metabolic changes were already present at that time, more advanced neurobiological changes may not yet have developed. Third, the study did not include direct brain measures, such as hippocampal inflammatory markers, synaptic plasticity, neurogenesis, insulin signalling or microglial activation. Fourth, some animals did not reach the TUNL acquisition criterion and were therefore not included in the later challenge phases, reducing the sample size for these analyses. Finally, full blinding was not feasible because the diets differed visibly, although automated data acquisition in the touchscreen chambers reduced observer-dependent scoring.

## Conclusion

Adolescent-onset HFHS diet exposure induced clear metabolic dysfunction and altered general behavioural patterns, but did not produce detectable impairments in object-based memory, TUNL acquisition, delayed spatial working memory or spatial pattern separation. These findings indicate that metabolic disruption after adolescent-onset obesogenic diet exposure does not necessarily coincide with measurable cognitive impairment. Future studies should determine whether longer exposure before the onset of behavioural testing, inclusion of female animals, or direct assessment of hippocampal and inflammatory mechanisms reveals conditions under which diet-induced metabolic dysfunction translates into cognitive decline.

## Acknowledgments

Funding: This research was funded by the European Union’s European Regional Development Fund (ERDF), project BriteN: Early Nutritional Interventions for Healthy Brain Development PROJ-00405, and Reckitt/Mead Johnson Nutrition, Nijmegen, The Netherlands. The preparation of the manuscript and Open Access publication costs were supported by ZonMw under grant/dossier number 01142042310007.

## References

1. Bach C, Souza JV, Oliveira CS, Mello RG, Maria-Ferreira D. Biochemical, molecular and behavioral changes induced by high-fat and high-sugar diets: a systematic review of non-clinical studies. Mol Neurobiol. 2026;63:524. doi:10.1007/s12035-026-05816-w

2. Freeman LR, Haley-Zitlin V, Rosenberger DS, Granholm AC. Damaging effects of a high-fat diet to the brain and cognition: a review of proposed mechanisms. Nutr Neurosci. 2014;17(6):241–251. doi:10.1179/1476830513Y.0000000092

3. Davidson TL, Hargrave SL, Swithers SE, Sample CH, Fu X, Kinzig KP, et al. Inter-relationships among diet, obesity and hippocampal-dependent cognitive function. Neuroscience. 2013;253:110–122. doi:10.1016/j.neuroscience.2013.08.044

4. Beilharz JE, Maniam J, Morris MJ. Diet-induced cognitive deficits: the role of fat and sugar, potential mechanisms and nutritional interventions. Nutrients. 2015;7(8):6719–6738. doi:10.3390/nu7085307

5. Kanoski SE, Davidson TL. Western diet consumption and cognitive impairment: links to hippocampal dysfunction and obesity. Physiol Behav. 2011;103(1):59–68. doi:10.1016/j.physbeh.2010.12.003

6. Reichelt AC. Examining adolescence as a sensitive period for high-fat, high-sugar diet exposure and cognition. Front Neurosci. 2019;13:1108. doi:10.3389/fnins.2019.01108

7. Spear LP. The adolescent brain and age-related behavioral manifestations. Neurosci Biobehav Rev. 2000;24(4):417–463. doi:10.1016/S0149-7634(00)00014-2

8. Del Olmo N, Ruiz-Gayo M. Influence of high-fat diets consumed during the juvenile period on hippocampal morphology and function. Front Cell Neurosci. 2018;12:439. doi:10.3389/fncel.2018.00439

9. Boitard C, Etchamendy N, Sauvant J, Aubert A, Tronel S, Marighetto A, et al. Juvenile, but not adult exposure to high-fat diet impairs relational memory and hippocampal neurogenesis in mice. Hippocampus. 2012;22(11):2095–2100. doi:10.1002/hipo.22032

10. Boitard C, Cavaroc A, Sauvant J, Aubert A, Castanon N, Layé S, et al. Impairment of hippocampal-dependent memory induced by juvenile high-fat diet intake is associated with enhanced hippocampal inflammation in rats. Brain Behav Immun. 2014;40:9–17. doi:10.1016/j.bbi.2014.03.005

11. Reichelt AC, Morris MJ, Westbrook RF. Daily access to sucrose impairs aspects of spatial memory tasks reliant on pattern separation and neural proliferation in rats. Learn Mem. 2016;23(7):386–390. doi:10.1101/lm.042416.116

12. Khazen T, Hatoum OA, Ferreira G, Maroun M. Acute exposure to a high-fat diet in juvenile male rats disrupts hippocampal-dependent memory and plasticity through glucocorticoids. Sci Rep. 2019;9:12270. doi:10.1038/s41598-019-48800-2

13. Noble EE, Olson CA, Davis E, Tsan L, Chen YW, Schade R, et al. Gut microbial taxa elevated by dietary sugar disrupt memory function. Transl Psychiatry. 2021;11:194. doi:10.1038/s41398-021-01309-7

14. Bakoyiannis I, Ducourneau EG, Santoyo-Zedillo M, Bosch-Bouju C, Pacheco-Lopez G, Coutureau E, et al. Obesogenic diet induces circuit-specific memory deficits in mice. eLife. 2024;13:e80388. doi:10.7554/eLife.80388

15. Deshpande NG, Saxena J, Pesaresi TG, Carrell CD, Ashby GB, Liao MK, et al. High fat diet alters gut microbiota but not spatial working memory in early middle-aged Sprague Dawley rats. PLoS One. 2019;14(5):e0217553. doi:10.1371/journal.pone.0217553

16. Leyh J, Winter K, Reinicke M, Ceglarek U, Bechmann I, Landmann J. Long-term diet-induced obesity does not lead to learning and memory impairment in adult mice. PLoS One. 2021;16(9):e0257921. doi:10.1371/journal.pone.0257921

17. Antunes M, Biala G. The novel object recognition memory: neurobiology, test procedure, and its modifications. Cogn Process. 2012;13(2):93–110. doi:10.1007/s10339-011-0430-z

18. Dix SL, Aggleton JP. Extending the spontaneous preference test of recognition: evidence of object-location and object-context recognition. Behav Brain Res. 1999;99(2):191–200. doi:10.1016/S0166-4328(98)00079-5

19. Talpos JC, McTighe SM, Dias R, Saksida LM, Bussey TJ. Trial-unique, delayed nonmatching-to-location: a novel, highly hippocampus-dependent automated touchscreen test of location memory and pattern separation. Neurobiol Learn Mem. 2010;94(3):341–352. doi:10.1016/j.nlm.2010.07.006

20. Oomen CA, Hvoslef-Eide M, Heath CJ, Mar AC, Horner AE, Bussey TJ, et al. The touchscreen operant platform for testing working memory and pattern separation in rats and mice. Nat Protoc. 2013;8(10):2006–2021. doi:10.1038/nprot.2013.124

21. Barnard IL, Onofrychuk TJ, McElroy DL, Howland JG. The touchscreen-based trial-unique, nonmatching-to-location task for rodents. Curr Protoc. 2021;1(10):e238. doi:10.1002/cpz1.238

22. Bennett D, Nakamura J, Vinnakota C, Sokolenko E, Nithianantharajah J, van den Buuse M, et al. Mouse behavior on the trial-unique nonmatching-to-location (TUNL) touchscreen task reflects a mixture of distinct working memory codes and response biases. J Neurosci. 2023;43(31):5693–5709. doi:10.1523/JNEUROSCI.2101-22.2023

23. Vinnakota C, Hudson MR, Ikeda K, Ide S, Mishina M, Sundram S, et al. Effects of NMDA receptor antagonists on working memory and gamma oscillations, and the mediating role of the GluN2D subunit. Neuropsychopharmacology. 2025;50:1938–1948. doi:10.1038/s41386-025-02129-9

24. Bassett AP, Bransom L, Mailman RB, Yang Y. Effect of functionally selective dopamine D1 receptor agonists on complex cognitive processes in a rodent touchscreen operant chamber task. eNeuro. 2026;13(5):ENEURO.0296-25.2026. doi:10.1523/ENEURO.0296-25.2026

25. Castillon CCM, Otsuka S, Armstrong JN, Contractor A. Subregional activity in the dentate gyrus is amplified during elevated cognitive demands. eLife. 2026;15:RP109611. doi:10.7554/eLife.109611.4

26. Percie du Sert N, Hurst V, Ahluwalia A, Alam S, Avey MT, Baker M, et al. The ARRIVE guidelines 2.0: updated guidelines for reporting animal research. PLoS Biol. 2020;18(7):e3000410. doi:10.1371/journal.pbio.3000410

27. Van de Weerd HA, Bulthuis RJA, Bergman AF, Schlingmann F, Tolboom J, Van Loo PLP, et al. Validation of a new system for the automatic registration of behaviour in mice and rats. Behav Processes. 2001;53(1–2):11–20. doi:10.1016/S0376-6357(00)00135-2

28. Ennaceur A, Delacour J. A new one-trial test for neurobiological studies of memory in rats. 1: Behavioral data. Behav Brain Res. 1988;31(1):47–59. doi:10.1016/0166-4328(88)90157-X

29. Bussey TJ, Padain TL, Skillings EA, Winters BD, Morton AJ, Saksida LM. The touchscreen cognitive testing method for rodents: how to get the best out of your rat. Learn Mem. 2008;15(7):516–523. doi:10.1101/lm.987808

30. Horner AE, Heath CJ, Hvoslef-Eide M, Kent BA, Kim CH, Nilsson SR, et al. The touchscreen operant platform for testing learning and memory in rats and mice. Nat Protoc. 2013;8(10):1961–1984. doi:10.1038/nprot.2013.122

31. Cardinal RN, Aitken MRF. Whisker: a client-server high-performance multimedia research control system. Behav Res Methods. 2010;42(4):1059–1071. doi:10.3758/BRM.42.4.1059

32. Spoelder M, Bright Y, Morrison MC, van Kempen V, de Groodt L, Begalli M, et al. Cognitive performance during the development of diabetes in the Zucker Diabetic Fatty rat. Cells. 2023;12(20):2463. doi:10.3390/cells12202463

33. Sachdeva AK, Dharavath RN, Chopra K. Time-response studies on development of cognitive deficits in an experimental model of insulin resistance. Clin Nutr. 2019;38(3):1447–1456. doi:10.1016/j.clnu.2018.05.018

34. Bae HR, Shin SK, Lee JY, Choi SS, Kwon EY. Chronological dynamics of neuroinflammatory responses in a high-fat diet mouse model. Int J Mol Sci. 2024;25(23):12834. doi:10.3390/ijms252312834

35. González Olmo BM, Bettes MN, DeMarsh JW, Zhao F, Askwith C, Barrientos RM. Short-term high-fat diet consumption impairs synaptic plasticity in the aged hippocampus via IL-1 signaling. NPJ Sci Food. 2023;7:35. doi:10.1038/s41538-023-00211-4

36. Zequeira S, Gazarov EA, Güvenli AA, Berthold EC, Senetra AS, Febo M, et al. Effects of cannabis smoke and oral Δ9THC on cognition in young adult and aged rats. Psychopharmacology. 2025;242:835–853. doi:10.1007/s00213-025-06754-6

37. Gopalan C, Niepoetter P, Butts-Wilmsmeyer C, Medavaka S, Ogle A, Daughrity S, et al. Comparison of intermittent fasting and voluntary wheel running on physical and cognitive abilities in high-fat diet-induced obese rats. PLoS One. 2023;18(12):e0293415. doi:10.1371/journal.pone.0293415

38. Barber TM, Kyrou I, Randeva HS, Weickert MO. Mechanisms of insulin resistance at the crossroad of obesity with associated metabolic abnormalities and cognitive dysfunction. Int J Mol Sci. 2021;22(2):546. doi:10.3390/ijms22020546

